# Particularity of “Universal resilience patterns in complex networks”

**DOI:** 10.1101/056218

**Authors:** Jean-Francois Arnoldi, Bart Haegeman, Tomas Revilla, Michel Loreau

**Affiliations:** Center for Biodiversity Theory and Modelling, CNRS, Moulis, France.; Biology Centre of the Academy of Sciences, Ceské Budejovice, Czech Republic.

## Abstract

In a recent *Letter to Nature*,Gao, Barzel and Barabási ^1^ describe an elegant procedure to reduce the dimensionality of complex dynamical networks, which they claim reveals “universal patterns of network resilience”, offering “ways to prevent the collapse of ecological, biological or economic systems, and guiding the design of technological systems resilient to both internal failures and environmental changes”. However, Gao et al restrict their attention to systems for which all interactions between nodes are mutualistic. Since antagonism is ubiquitous in natural and social networks, we clarify why this stringent hypothesis is necessary and what happens when it is relaxed. By analyzing broad classes of competitive and predator-prey networks we provide novel insights into the underlying mechanisms at work in Gao et al’s theory, and novel predictions for dynamical systems that are not purely mutualistic.

---

Following ref. 1, we consider a network of *N* nodes whose activities *x_i_* are governed by

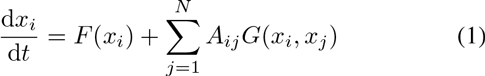

F(*x_i_*) describes self-dynamics of node *i* while *G*(*x_i_, x_j_*) represents the effect of node *j* on node *i*. *A* is a matrix of weighted links, encoding network structure. At the core of Gao et al’s^1^ procedure is a *mean-field* approximation, replacing all interacting partners of any focal node by replicates of a single partner *x*_eff_ = **1**^⊤^ ***Ax***/**1**^⊤^ ***A*1**, where ***x*** = (*x_1_,…,x_N_*)^⊤^ and **1** = (1,…,1)^⊤^. In a second step the dynamics of x_eff_ are reduced to

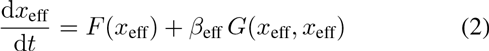

where *β*_eff_ = **1**^⊤^*A*^2^**1**/**1**^⊤^*A***1**. In this approximation, the state of the effective node determines the state of the entire system. In the following, we assume the network to be large, of maximal connectance and that self-regulation is homogeneous amongst nodes. *A priori*, this is an ideal setting for a mean-field approximation. We begin with linear interactions, corresponding to *F*(*x_i_*) = − *x_i_* and *G*(*x_i_,x_j_*) = *gx_j_*. Since the seminal work of May^2^, linear systems have served as a benchmark for the study of ecological stability^2,3^. In both mutualisitic (*g* = +1) and competitive (*g* = −1) cases, Gao et al’s procedure is now exact when all interaction rates *A_ij_* are equal. In compact form, the activity of nodes follow *d**x**/dt* = *M **x*** where 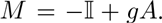

The stability criterion is that the dominant eigenvalue λ_true_ of the matrix *M* has negative real part. The effective node should follow *dx*_eff_/*dt* = λ_eff_*x*_eff_ where λ_eff_ = − 1 + *gβ*_eff_, so that the approximate stability criterion is that λ_eff_ is negative. We draw interaction rates *A_ij_* > 0 as independent realizations of the random variable |*X*| where 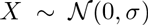, taking *σ* as a measure of mean interaction strength. In the first panel of Fig. 1, we generated mutualistic networks for various values of *σ* comparing true and predicted stability. We observe a perfect correspondence between the two. This can be explained by the fact that, in those large randomly assembled networks, the dominant eigenvalue of *A* is well approximated by *β*_eff_ while the associated eigenvector approaches **1**, a mathematical expression of the mean-field approximation^3^. When *g* is positive this implies that λ_eff_ is a good approximation of the dominant eigenvalue of *M*. The network is thus stable if and only if the approximate system is stable, in agreement with the examples of ref. 1. Importantly, *x_eff_* approaches the mean activity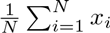–in an ecological context, mean biomass; projection of ***x*** on the direction spanned by **1**, the slowest, least stable, direction of the dynamics. The situation changes dramatically when considering competitive interactions. Indeed we now have that λ_eff_ is a good approximation of the *smallest* eigenvalue of *M*, which does not determine stability. On the second panel of Fig. 1, we generated competitive communities along a gradient of interaction strength. A sharp divergence between true and approximated network stability is observed. In fact, for a large number *N* of nodes, the dominant eigenvalue of *M* approaches^3^

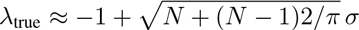

indicating a loss of stability as *N* and/or σ increase, whereas

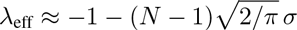

gives the opposite trend. Mean activity is now the fastest, most stable, direction of the dynamics, while differences between individual activities can grow uncontrollably. Although the state of the effective node should be approximated with reasonable precision by Gao et al’s reduction, it is not sufficient to predict the state of the entire network. The negative impact of one node on the other has a positive impact on other competing nodes, making unpredictable the net feedback experienced by a focal node. The effect of neighboring nodes cannot be reduced to an effective competitor. This last point is particularly obvious for networks of predator and prey, whose interactions are both positive *and* negative. If for any interacting pairs we decide at random who is prey and who is predator, as the number of nodes *N* grows, λ_true_ follows a universal trend^3^:

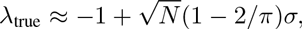

where as λ_eff_ is ill-defined as the denominator of *β*_eff_ has now zero expected value. In other words, the net balance between positive and negative feedbacks experienced by a focal node can switch unpredictably by the addition or removal of a single species, preventing the mean-field approximation to hold. This explains the dispersion of predicted stability in the last panel of Fig. 1.

**Figure 1:**
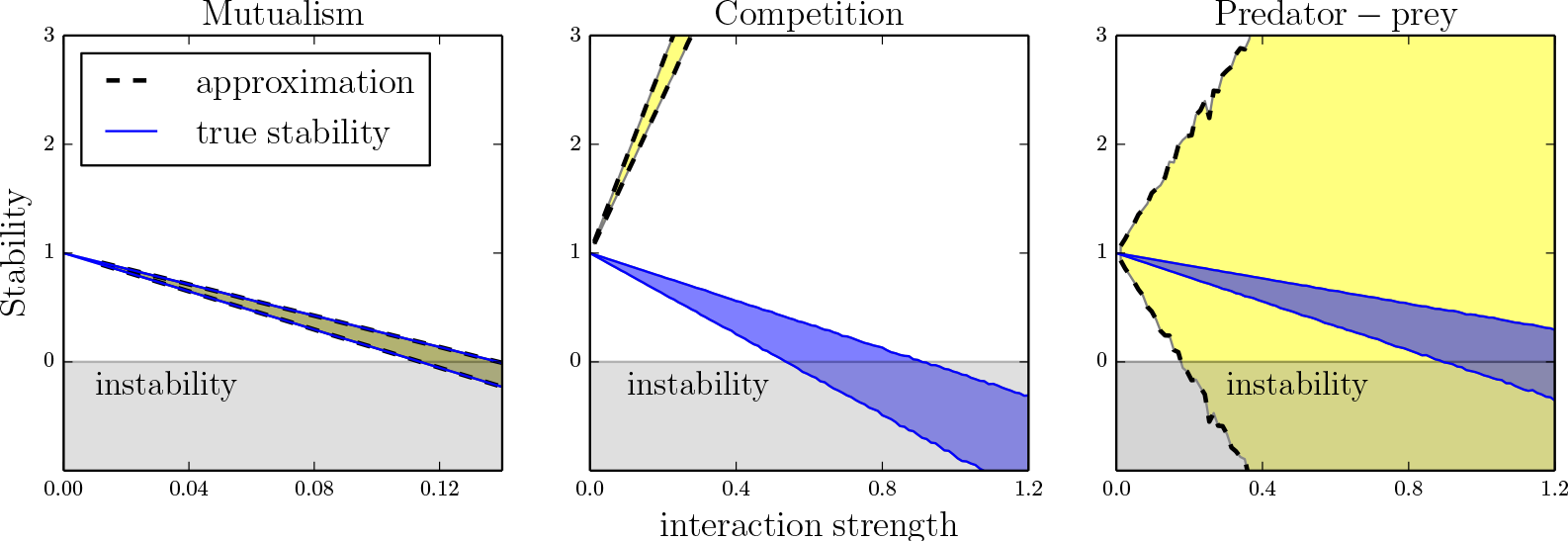
Stability of linear systems. Along a gradient of mean interaction strength, we randomly generated 10000 fully connected networks composed of *N* = 10 dynamical nodes, representing three types of ecological communities near equilibrium. The region shaded in blue represent the 10 − 90 percentiles of realized stability −ℜ(λ_true_) (unstable if negative). In yellow is the equivalent range for predicted stability −λ_eff_.

When interactions are non-linear, several dynamical states can coexist, ranging from equilibriums to chaos, and stability is defined relatively to alternate, undesirable states. For mutualistic systems, Gao et al provide compelling examples indicating that their theory reveals reliable stability patterns. The above analysis of linear systems allows us to draw a qualitative understanding of why this can be true: Suppose the interaction matrix *A* has an eigenvalue β_eff_ associated to **1**. Any non-linear dynamics of the form (1) will then preserve the direction spanned by **1**, following a linear trajectory *φ*(*t*) = *x*_eff_(*t*)**1** with *x*_eff_ solution of the reduced system (2). In mutualistic systems, if the reduced system converges to an equilibrium *x*̂_eff_, *φ*(*t*) describes the trajectory along the direction of slowest approach to the full steady-state x̂_eff_**1**/‖**1**‖. A change in stability of a steady state will automatically be seen on the reduced system. In complex mutualistic networks, **1** will typically be a good approximation of the eigenvector associated to *β*_eff_, so that we can expect the described behavior to hold, at least qualitatively. If interactions are purely competitive all of the above remains valid, except that the trajectory ***φ***(*t*) will not be the one of slowest approach suggesting that instabilities may go undetected. As an example, let us revisit the case of competitive systems, of the form of generalized Lotka-Volterra communities, for which *F*(*x_i_*) = *x_i_*(1 − *x_i_*) and *G*(*x_i_, x_j_*) = *gx_i_x_j_*, with *g* = −1. Once reduced by Gao et al’s procedure, the effective node follows logistic growth

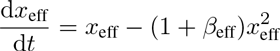

and approximates, as in the linear case, the system’s mean activity. Because *β*_eff_ is positive, we see that the reduction predicts a unique effective stable equilibrium *x*̂_eff_ = 1/(1+*β*_eff_). Validity of Gao et al’s reduction in this setting would imply that such systems present, at most, various alternate stable states corresponding to the same mean activity. In Fig. 2, with similar rules as in the linear case, we generated a community composed of *N* = 23 competing species. Along a gradient of interaction strength, alternate stable states coexist, corresponding to different assemblages of persisting species. Remarkably, mean biomass *x*_eff_ is similar for different stable states and well predicted by Gao et al’s approximation. This, however, also implies that the reduction is not informative in terms of persistence or abundances of species, a limitation worth pointing out in view of application to conservation issues. Furthermore, stable states can coexist with limit cycles or chaos, which are undetected by the reduced system. In Fig. 2, one equilibrium undergoes a Hopf bifurcation to a limit cycle presenting large oscillations. At this point, the possible transition from a remaining equilibrium to undesired oscillations is not accounted for.

Let us now briefly comment the more general case of a mixture of interactions types. Starting from a purely mutualistic context, if antagonism is gradually allowed in the system, the eigenvector associated to the dominant eigenvalue of A will soon depart from the homogeneous vector **1** so that ***φ***(*t*) = *x*_eff_(*t*)**1** will not be a solution anymore (even if *β*_eff_ remains a good approximation of the dominant eigenvalue of the interaction matrix *A*, even-though it would suffice to predict the qualitative state of a linear system). Due to non-linearities, there will not be any linear direction preserved by the dynamics along which it reduces to the simplified system (2). All in all, this makes it difficult to foresee the ability of Gao et al’s procedure to predict qualitative stability patterns. In conclusion, it is unclear if a statistical theory of complex networks can exist. Gao et al show that reliable emerging stability properties can be expected when interactions are purely mutualistic. We explained why these properties are specific to mutualism and should not be expected to hold in networks for which antagonistic interactions cannot be neglected. This limitation is important to keep in mind before applying this method to real-world systems, in which antagonism is ubiquitous and often a dominant form of interaction.

**Figure 2:**
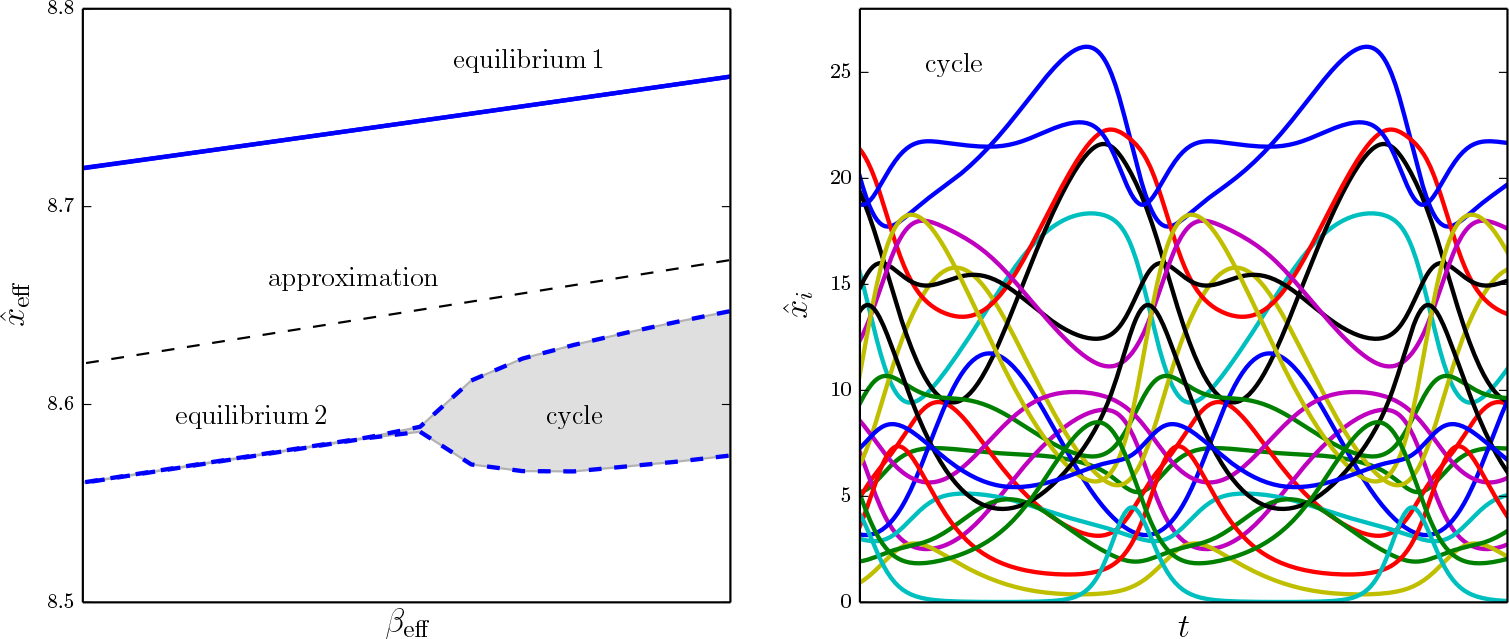
Detail of bifurcation diagram of a competitive Lotka-Volterra system. We randomly generated a community composed of *N* = 23 species. Along a gradient of interaction strength, alternate stable state coexist, corresponding to different assemblages of persisting species (first panel, equilibrium 1 and 2). Associated effective variable *x*_eff_ is similar for all and well predicted by the approximation of Gao et al. Equilibrium 2 undergoes a Hopf bifurcation to a limit cycle (first panel, shaded gray region) presenting large oscillations (right panel).

## Acknowledgements

The authors would like to thank Baruch Barzel and coauthors for taking the time to kindly respond to our comments on their work. This work is supported by the TULIP Laboratory of Excellence (ANR-10-LABX-41), the AnaEE-France project (ANR-11-INBS-0001) and by the BIOSTASES Advanced Grant, funded by the European research council under the European Union’s Horizon 2020 research and innovation programme (grant agreement No 666971).

